# Immediate and Long-Term Electrophysiological Biomarkers of Antidepressant Behavioral Effects After Subanesthetic Ketamine and Medial Prefrontal Cortex Deep Brain Stimulation Treatment

**DOI:** 10.1101/2023.10.13.562089

**Authors:** Matthew Bergosh, Sasha Medvidovic, Nancy Zepeda, Lindsey Crown, Jennifer Ipe, Lauren Debattista, Luis Romero, Eimon Amjadi, Tian Lam, Erik Hakopian, Wooseong Choi, Kevin Wu, Yu Tung Jack Lo, Darrin Jason Lee

## Abstract

1

Ketamine and medial prefrontal cortex (mPFC) deep brain stimulation (DBS) are emerging depression treatments, however their mechanisms and efficacy biomarkers have not been established. This study investigates immediate and long-term mPFC electrophysiological correlates of their antidepressant effects, and explores the effects of combinatorial treatment. Local field potential electrodes were implanted into the mPFC of male Sprague Dawley rats, then corticosterone was administered for 21 days, except in the control group (n = 8). In the last week of depression induction, the remaining groups received mPFC DBS (n = 8), ketamine (n = 9), both (n = 9), or neither (n = 8). All groups then performed a behavioral assay. We found that DBS treatment immediately suppressed relative low gamma power and improved cognition in the long-term, which correlated with rescued apathy-like behavior in the Groom Test (GT). In contrast, ketamine treatment suppressed sample entropy in the long-term, which correlated with its immediate decrease in sample entropy as well as rescued GT behavior. In the combinatorial group these changes were inhibited, which correlated with a failure to fully rescue GT behavior. Despair-like behavior in the Forced Swim Test was rescued by all three treatments, which correlated with increased relative high frequency oscillation (HFO) power immediately after ketamine and combinatorial treatment. Our findings support the utility of low gamma and HFO power, sample entropy, and cognitive tests in the search for mechanisms and biomarkers of depression and novel treatments.

**Significance Statement:** There is growing interest in psychedelic pharmacological agents, such as ketamine, and direct neurostimulation techniques, like deep brain stimulation, as treatments for depression. However, their mechanisms are poorly understood, particularly those underlying their sustained benefits. Our study addresses this by correlating behavioral rescue with immediate and long-term changes in the function of the medial prefrontal cortex. These antidepressant correlates may represent therapeutically relevant functional changes, and therefore also have potential as biomarkers. Clinically, such biomarkers enable predicting and monitoring of treatment efficacy, and thereby contribute to the personalization of care. In research, utilizing these translatable biomarkers enhances the etiological validity of disease models, and enables the development of more targeted, next-generation treatment strategies.

## 3. Introduction

Depression is the most common psychiatric condition globally, affecting an estimated 246 million people in 2020 (Santomauro et al., 2021), with an increasing prevalence in recent years (Goodwin et al., 2022). Yet current medications are only effective for 30-40% of patients, and require weeks to achieve a therapeutic effect, so there is a growing unmet need for more effective treatments (Blackburn, 2019). Recently, there has been mounting preclinical and clinical evidence supporting emerging treatments such as deep brain stimulation (DBS) of the medial prefrontal cortex (mPFC) (Dandekar et al., 2018), as well as subanesthetic doses of ketamine (Bobo et al., 2016).

However, their therapeutic mechanisms are not fully understood. Oscillatory activity in local field potential (LFP) and electroencephalogram recordings has been used to study the neurophysiological effects of both of these treatment modalities in depression. Nearly every canonical frequency band has been implicated in depression or remission, especially within the mPFC. Changes in the power of frequency bands here can act as biomarkers for treatment efficacy (Fitzgerald & Watson, 2018), as well as treatment mechanisms, as they reflect changes in the function of the mPFC. Similarly, changes in measures of signal complexity and irregularity, such as sample entropy, can provide insight into the functional neural changes that underlie depression and the action of treatments (Lin et al., 2020; Wang et al., 2018).

These mPFC activity changes, and correlations with behavior, after ketamine and mPFC DBS have never been investigated within a single study, so it is difficult to determine whether apparent electrophysiological and behavioral differences merely result from differing methodology. Furthermore, no preclinical studies have attempted to connect immediate mPFC activity changes after DBS or ketamine treatment to long-term (>24 hours) changes in mPFC activity, or associate long-term changes here with sustained antidepressant effects. Additionally, no studies have attempted to combine these treatment modalities, despite evidence of overlapping mechanisms, such as beta and gamma power modulation (Fitzgerald & Watson, 2018; Jia et al., 2019; Sun et al., 2015) or synaptic plasticity and potentiation (Bambico et al., 2015; Jett et al., 2015; Sumner et al., 2020; Sun et al., 2015).

Therefore, in the present study, we investigated electrophysiological changes in the rat mPFC homolog, and behavioral effects of the well-established chronic corticosterone (CORT) preclinical depression model, as well as repeated treatment by mPFC DBS, ketamine, or both. We hypothesized that the combination of treatments would induce synergistic behavioral and electrophysiological antidepressant effects. LFP recordings immediately, one day, and one week after treatment offset were compared to baseline recordings and between groups, and then correlated with depression- or remission-like performance in a behavioral assay. We hypothesized that changes in the activity of the mPFC would parallel, and possibly drive, the antidepressant action of ketamine and mPFC DBS, and therefore could act as etiological and therapeutic biomarkers.

## 4. Methods

### 4.1 Animal numbers, housing, and groups

42 male 6-8 week old (350 ± 50g) Sprague-Dawley rats (Charles River Laboratories, Wilmington, MA) were obtained. The animals acclimated to the facilities for 1 week before experimental procedures began. Animals had access to food and water ad libitum, except for 24 hours preceding the Sucrose Preference Tests. Following the electrode placement surgery, the animals were single housed for the remainder of the experiment, to protect the electrode implant from cagemates. The animals were kept in a controlled environment at 23 ± 1◦C, 55–65% relative humidity, and a fixed 12 hour light/dark cycle with lights on at 0600. All procedures performed were approved by the [Author University] Institutional Animal Care and Use Committee. Animals were weighed and observed daily for the duration of the experiment. The animals were randomly placed into one of five groups: control (n = 8), untreated (n = 8), ketamine (n = 9), DBS (n = 8), and DBS+ketamine (n = 9).

### 4.2 Electrode implantation surgery

An overview of the experimental design is given in Figure 1A. After acclimation, the animals underwent electrode implantation surgery. Under general anesthesia (2-4% isoflurane in O_2_ carrier), a midline incision was made in the scalp, starting just behind the eyes and finishing just behind the ears. Using a digital stereotaxic frame, twisted bipolar tungsten electrodes (P1 Technologies, Roanoke County, Virginia) were inserted into the mPFC, including the prelimbic cortex and primary cingulate cortex (ML: 0.6 mm relative to bregma, AP: 2.5 mm, DV: 4.0 mm) (Fig. 1B) and lateral visual cortex (ML: 5.5 mm, AP: −6.0 mm, DV: 2.5 mm) (Fig. 1C) (Paxinos & Watson, 2007). Six partial thickness holes were drilled into the skull, and six stainless steel autoclave-sterilized small anchoring screws (#0-80) were threaded into the skull. The electrodes were then anchored in place using methyl methacrylate.

**Figure 1.**
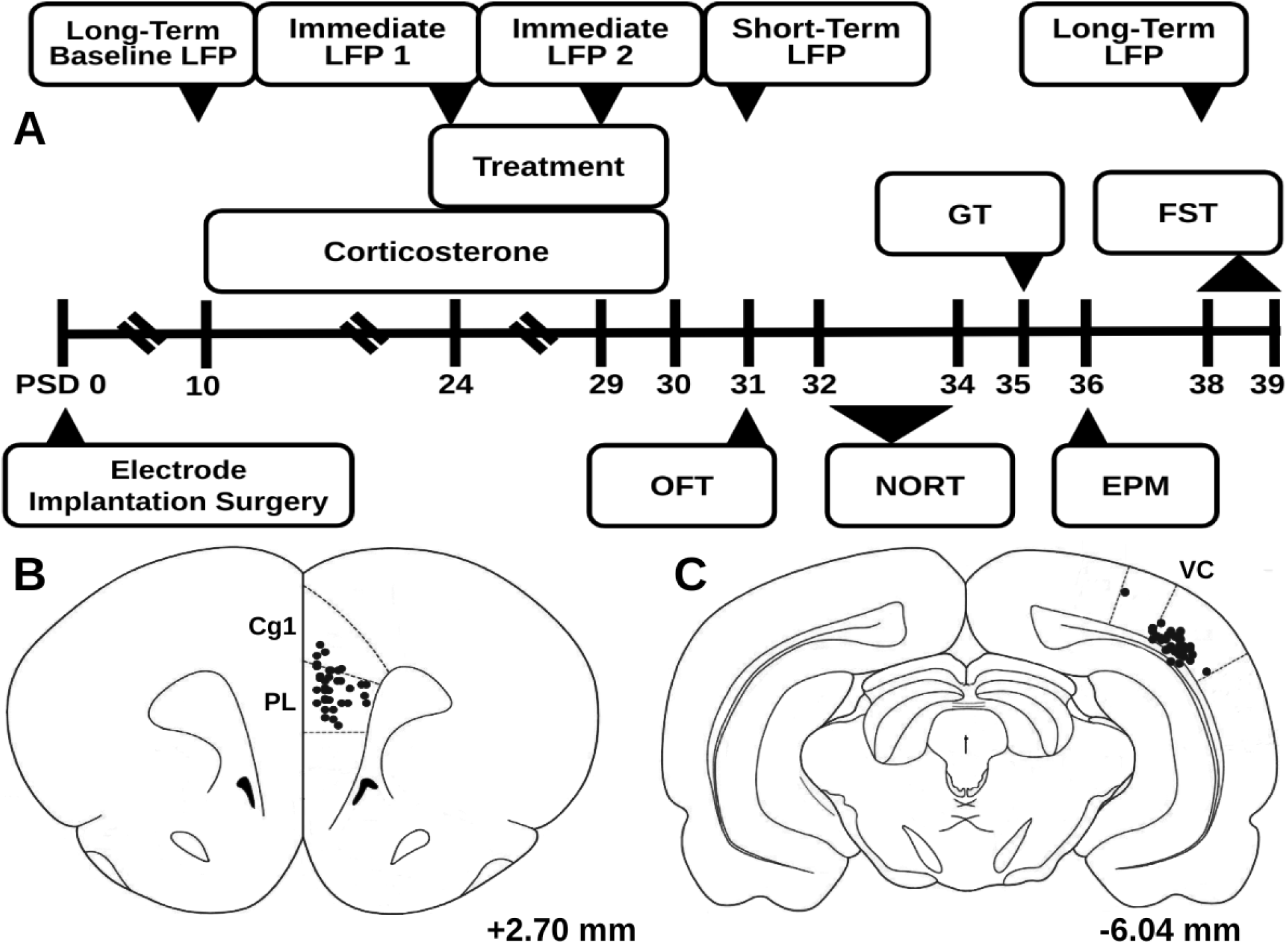
Experiment overview. A, Timeline of experiment. Below, electrode tip locations are marked on a modified atlas with black dots. B, The region of interest, the medial prefrontal cortex, including the primary cingulate cortex (Cg1) and the prelimbic cortex (PL). This region was the target of the deep brain stimulation treatment, and studied through local field potential (LFP) recordings throughout the experiment. C, The structure referenced from during LFP recordings, the lateral visual cortex (VC). Atlas illustrations adapted from Paxinos & Watson, 2007, and the given plane coordinates are referenced from bregma. (PSD = post surgery day, OFT = Open Field Test, NORT = Novel Object Recognition Test, GT = Groom Test, EPM = Elevated Plus Maze, FST = Forced Swim Test)

### 4.3 Pharmacology and stimulation

#### 4.3.1 Corticosterone depression model

On post surgery days (PSD) 10-30, 40 mg/kg CORT (C2505, Sigma-Aldrich, Milwaukee, WI) was subcutaneously administered, except in the control group which received 5% DMSO/saline vehicle. Previous studies have demonstrated that this dose and duration induces depressive-like symptoms, such as increases in anhedonia, anxiety, and despair, decreases in evoked grooming behavior, impairments in declarative memory, and weight disturbances (Sterner & Kalynchuk, 2010). Animals received the CORT dose after LFP recordings, treatment administrations, and behavioral tasks were completed for the day.

#### 4.3.2 Ketamine treatment

On PSD 24-30, 15 mg/kg ketamine hydrochloride (K2753, Sigma-Aldrich, Milwaukee, WI) were intraperitoneally administered to the ketamine and DBS+ketamine groups. This dose and duration was chosen based on previous studies (Garcia et al., 2008), which show a rescue of several depressive-like symptoms listed in the CORT section. Saline vehicle was administered to the control, untreated, and DBS-only groups.

#### 4.3.3 Deep brain stimulation treatment

On PSD 24-30 the DBS and DBS+ketamine animals’ implanted mPFC electrode was connected to an isolated pulse stimulator (STG 4008, Warner Instruments LLC, Hamden, CT) via a twisted wire cable. Continuous electrical stimulation was delivered using the following stimulation parameters: 100 microsecond square wave pulses at 130 Hz and current of 80 μA for 30 minutes. These parameters were selected based on literature (Hamani et al., 2012; Hamani & Nóbrega, 2010), which demonstrated a rescue of several depressive-like symptoms listed in the CORT section. Animals in the DBS+ketamine group were administered the ketamine dose immediately before stimulation began. Sham stimulation was administered to the control, untreated, and ketamine-only groups.

### 4.4 Behavioral tests and weight

#### 4.4.1 Open Field Test (OFT)

Non-specific changes in locomotor activity after corticosterone, ketamine, and DBS (Karatsoreos et al., 2010; Papp et al., 2022; Parise et al., 2013) can confound behavioral and electrophysiological results, therefore we measured average speed in the Open Field Test on PSD 31. Animals were placed in an open, square arena (50 x 50 cm) and allowed to explore for 10 minutes. The average speed of the animal was measured using the video tracking software TopScan Lite (Clever Sys Inc., Reston, VA). Similarly, non-specific changes in grooming behavior can confound the Groom Test (Planchez et al., 2019), therefore the number of seconds spent spontaneously grooming during the OFT were evaluated.

#### 4.4.2 Novel Object Recognition (NORT)

Cognitive dysfunction, including deficits in memory, occur in depression (American Psychiatric Association, 2013). To measure these, we employed the Novel Object Recognition Test (NORT). In this test, a higher proportion of time spent with a novel object, compared to a familiar object, is interpreted as representing functional object recognition memory (Antunes & Biala, 2012). On PSD 31, the animals were habituated to the test arena during the OFT for 10 minutes. On PSD 32, they explored the arena which contained two identical objects: Object A, a black Monster Energy can (Monster Energy Company, Corona, CA), in the position it would occupy in all trials; and Object A’, in the position the novel objects would occupy. On PSD 33 the arena contained Object A; and a novel object, Object B, a white Monster Energy can. On PSD 34 the arena contained Object A; and a new novel object, Object C, a Powerade bottle (The Coca-Cola Company, Atlanta, GA). Video recordings were analyzed of each 5 minute trial on PSD 32-34 to quantify the number of seconds spent by the animal exploring each object. The Discrimination Index (DI) of each trial was calculated by subtracting the time each animal spent exploring the familiar object (A) from the time spent with the novel object (A’,B,C), and dividing by the total time exploring both objects. DI results of both test days (PSD 33 and 34) of an animal were averaged, then the DI of the familiarization day (PSD 32) was subtracted. This difference was normalized by dividing it by the familiarization day DI and multiplying by 100 to generate a percent change that was averaged across animals in a group.

#### 4.4.3 Groom Test (GT)

As healthy rodents groom themselves thoroughly in response to soiling of their coats, reduced evoked grooming in the Groom Test (GT) has been interpreted to represent apathy, a core symptom of depression (Kennedy, 2008; Planchez et al., 2019). The animals performed the GT on PSD 35, their dorsoposterior coats were sprayed with a 10% sucrose solution, and were observed for 10 minutes in the OFT/NORT arena. Video recordings were evaluated for the number of seconds spent grooming.

#### 4.4.4 Elevated Plus Maze (EPM)

Anxiety is highly comorbid with depression (American Psychiatric Association, 2013), and related to the mechanism of the chronic CORT model (Sterner & Kalynchuk, 2010). Therefore, on PSD 36, the Elevated Plus Maze (EPM) was used to measure anxiety, where a higher proportion of time in the innately fear-inducing “open” arms is interpreted as less baseline anxiety (Planchez et al., 2019). The EPM is a plus-shaped maze, with two opposing “closed” arms with sidewalls (40 cm tall), and two opposing “open” arms with no sidewalls. The lanes created by two opposing arms are 110 cm long, 10 cm wide, and 62 cm high. The animals were placed at the center of the maze and explored for 5 minutes. Video recordings were analyzed for the number of seconds spent in the open arms, which was divided by the total seconds of the trial, to calculate the percent of time spent in the open arms.

#### 4.4.5 Forced Swim Test (FST)

Despair, the second core symptom of depression beside apathy (Kennedy, 2008), is measured in rodents via the Forced Swim Test (FST), where increased immobile time in the test trial putatively represents despair (Planchez et al., 2019). On PSD 38, the animals were habituated to the FST testing chamber (glass cylindrical tub 50 cm tall, 25 cm diameter, filled with 21°C water 40 cm high) for 10 minutes. On PSD 39, the animals were tested for 5 minutes. A video recording of these five minutes was analyzed to determine the number of seconds spent immobile. After each trial the animals were placed in a heated chamber, dried, and monitored for full recovery.

#### 4.4.6 Weight

Weight disturbances are a common symptom in depression (American Psychiatric Association, 2013). Furthermore, similar to locomotor activity, non-specific changes in weight can confound behavioral results. Therefore, weights were measured at the “pre” CORT/treatment timepoint PSD 8-10, as well as the “post” timepoints PSD 30-32 and 37-39. Relative weights were calculated by dividing an animal’s weight at the “post” timepoints by the “pre” timepoint. The average of three days was used due to high variability in weight caused by the SPT fasts.

### 4.5 Local field potential recordings, spectral power and sample entropy calculations

To provide a baseline from which to track the short- and long-term oscillatory changes induced by chronic CORT and treatment, a 10 minute local field potential (LFP) recording of the mPFC was taken on PSD 10, prior to the first CORT dose, using a Cheetah Digital Lynx SX Data Acquisition System (Neuralynx, Bozeman, MT). These recordings were referenced against an arbitrary cortical region, the lateral secondary visual cortex (V2L). To standardize recording quality, segments at least 3 seconds long and containing less than 2% noise were analyzed. A power spectral density plot was generated using the pwelch() function in MATLAB (MathWorks, Natick, MA). To investigate long-term changes following the last CORT and/or treatment administration on PSD 30, 10 minute LFP recordings on PSD 31 and PSD 38 were taken. To investigate the immediate oscillatory effects induced by treatment administration, 10 minute baseline and 20 minute post-treatment recordings were taken on PSD 24 and PSD 29. All recordings occurred in a designated arena (30 cm x 46 cm x 19 cm).

Power values for canonical bands delta (1-3 Hz), theta (4-8 Hz), alpha (9-14 Hz), beta (15-30 Hz), low gamma (30-50 Hz), high gamma (50-130 Hz), and HFO (130-180 Hz) were obtained by applying a 12th order butterworth bandpass filter and taking the absolute value of the Hilbert transformed data. Similar notch filters were used in pre-processing to remove 60 Hz, 120 Hz, and 180 Hz noise. Relative power was obtained by dividing the power of each frequency band by the total power of all bands. Additionally, sample entropy, a measure of signal irregularity, was calculated using the sampen() function in MATLAB (Víctor Martínez-Cagiga, 2018), with r = .15 and m = 2, based on recommendations by previous studies (Molina-Picó et al., 2011). Percent change in sample entropy and the relative power of each frequency band was calculated by subtracting the value from the baseline recording (Pre-CORT: PSD 10; Pre-Treatment: PSD 24 and 29) from the value in the recording of interest (Post-CORT: PSD 31 and 38; Post-Treatment: PSD 24 and 29), dividing by the value of the baseline recording, and multiplying by 100. This process was also applied to the power spectral density plots, for the averaged changes on PSD 24 and 29, as well as those that occurred by PSD 38.

### 4.6 Statistical Analysis

Statistical analyses were performed using R (R Core Team, 2021). T-tests and/or ANOVA were used to compare between groups, while one-sample tests revealed if a group changed significantly relative to their baseline (mu = 0). Standard parametric tests (one- or two sample t-test, ANOVA, Pearson’s r) were used when assumptions of normality and homoscedasticity were met, while non-parametric tests (one- or two-sample Wilcoxon rank sum test, aligned rank transformed ANOVA, Spearman’s ρ) were used otherwise. False Discovery Rate corrections to p-values were used to control for familywise error. For all tests, alpha was set to 0.05 (two-tailed).

### 4.7 Histology

On PSD 39, rats were euthanized by anesthesia (Isoflurane) and were transcardially perfused with 100 ml of 0.1 M sodium phosphate buffer saline (PBS, pH −7.4), followed by 50 ml of 4% paraformaldehyde (pH 7.4). Brains were extracted and stored in 4% paraformaldehyde at 4 °C. Serial coronal sections were cut at 100-μm thickness with a vibratome (Leica VT 1200; Leica Biosystems, Buffalo Grove, IL) starting at +3.8 mm Bregma and ending at −6.50 mm Bregma. Sections in the vicinity of electrodes were mounted onto 0.1% gelatin-subbed slides and stained with NeuroTrace 530/615 Red Fluorescent Nissl Stain (N21482, ThermoFisher Scientific, Waltham, MA, USA) to confirm proper placement (Fig. 1B, C).

## 5. Results

### 5.1 Behavioral changes after chronic CORT and treatment

#### 5.1.1 Chronic CORT administration induced apathy-like symptoms in the GT, which was fully rescued by separate chronic DBS or ketamine treatment, but not by the combinatorial treatment

The results of the behavioral assay are shown in Figure 2. During the Groom Test (Fig. 2A), the untreated group (M = 56.6 s, SD = 42.4) spent significantly less time grooming than the control group (M = 133.9 s, SD = 51.1, p = .028), indicating apathy-like symptoms had been induced in the untreated group. The groups that received DBS (M = 155.6 s, SD = 82.1) or ketamine (M = 137.0 s, SD = 66.7) treatment separately spent significantly more time grooming than the untreated group (DBS: p = .028; ketamine: p = .028), and an equal amount to the control group, representing a complete rescue of apathy-like symptoms. The group that received both treatments concurrently (M = 105.2 s, SD = 82.8) did not significantly differ from the untreated group, but also did not differ from the control group, suggesting apathy-like symptoms were only partially rescued in this group. In addition, non-specific alterations in grooming activity can confound the GT, however, spontaneous grooming behavior in the OFT did not differ between groups (Fig. 2E).

**Figure 2.**
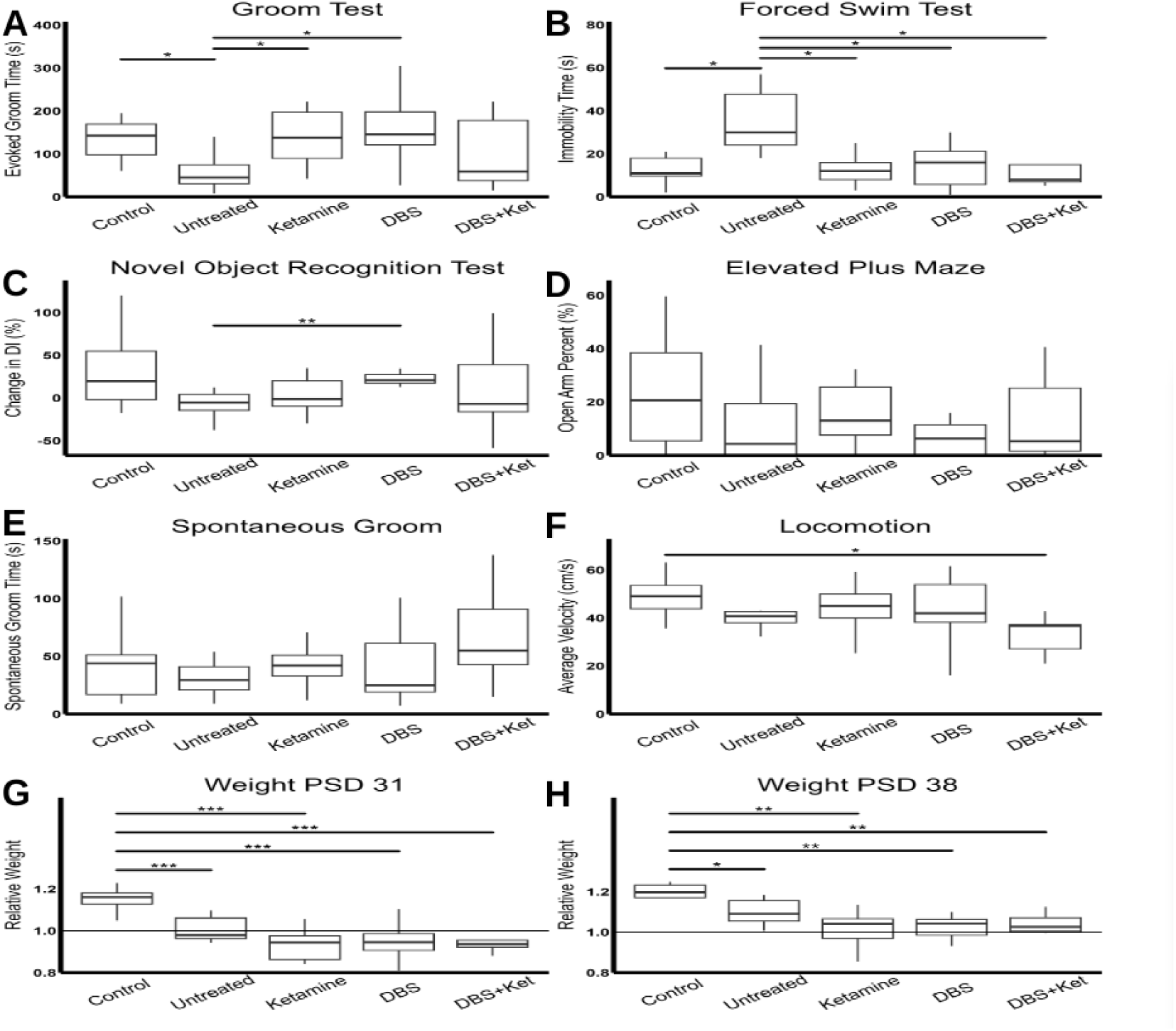
Behavioral and weight changes. A, CORT induced apathy-like symptoms in the untreated group by reducing evoked groom time in the GT, which ketamine and DBS separately, but not combined, rescued. B, CORT induced despair-like symptoms in the untreated group in the FST, which was rescued by ketamine, DBS, and their combination. C, CORT did not induce deficits in recognition memory in the NORT, however recognition memory was improved in the DBS group in comparison to the untreated group. CORT and treatments did not induce changes in an anxiety measure, the EPM (D), as well as an important control to the GT, spontaneous grooming in the OFT (E). F, CORT did not alter average velocity in the OFT, nor did treatment with ketamine or DBS separately, however the combination treatment reduced it. CORT induced depression-like deficits in weight gain that were not rescued at the short-(G) or long-term (H) timepoints by any treatment. (*: p < .05; **: p < .01; ***: p < .001; CORT = corticosterone, GT = Groom Test, DBS = Deep Brain Stimulation, FST = Forced Swim Test, NORT = Novel Object Recognition Test, EPM = Elevated Plus Maze, OFT = Open Field Test)

#### 5.1.2 Chronic CORT administration induced despair-like behavior in the FST, which was rescued by chronic DBS, ketamine, and the combinatorial treatment

Compared to the control group (M = 14.4 s, SD = 9.7), the untreated group (M = 35.3 s, SD = 15.5) spent more time immobile in the FST test (p = .012, Fig. 2B), indicating despair-like symptoms had been induced in the untreated group. This was rescued by all three treatments, who spent less time immobile than the untreated group (DBS: M = 14.6 s, SD = 9.9, p = .022; ketamine: M = 15.6 s, SD = 13.6, p = .022; DBS+Ket: M = 15.1 s, SD = 13.1, p = .022, all FDR corrected) and did not differ from the control group, indicating their performance had been normalized.

#### 5.1.3 Anxiety (EPM) and object recognition memory (NORT) measures were unaltered by chronic CORT, however DBS improved NORT performance

In the NORT (Fig. 2C), the percent change in Discrimination Index was greater in the DBS group (M = 28.8%, SD = 28.7) than the untreated group (M = −7.7%, SD = 17.1, p < .01). This indicates an improvement in object memory, however, as there was no deficit in the untreated group compared to the control group (M = 31.1%, SD = 46.9, p>0.05), this does not represent a “rescue” of memory function. There were no differences in open arm time percentage in the EPM between the untreated and control or treatment groups, indicating that, as has been previously reported (Bertholomey et al., 2022), comorbid anxiety-like symptoms were not induced (Fig. 2D).

#### 5.1.5 Chronic CORT induced weight disruptions that were not rescued by any treatment, and long-term locomotor activity was only altered in the DBS+ketamine group

The average relative weight (Fig. 2G, H) in the untreated group was less than in the control on PSD 31 (Control: M = 1.15, SD = .054; Untreated: M = 1.00, SD = .060; p < .001) and PSD 38 (Control: M = 1.19, SD = .060; Untreated: M = 1.10, SD = .07; p = .017), indicating depression-like weight disturbances had been induced. The three treatment groups did not differ from the untreated group, and had lower relative weights than the control group on PSD 31 (DBS: M = .95, SD = .087, p < .001; Ket: M = .94, SD = .078, p < .0001; DBS+Ket: M = .95, SD = .076, p < .0001) and PSD 38 (DBS: M = 1.04, SD = .082, p < .001; Ket: M = 1.02, SD = .090, p < .001; DBS+Ket: M = 1.04, SD = .080, p < .001). Therefore none of the treatment strategies tested rescued these disruptions in weight. No groups differed from the untreated group in average velocity in the OFT (Fig. 2F), though the DBS+ketamine group (M = 31.1 cm/s, SD = 12.6) had lower average velocities than the control group (M = 48.9 cm/s, SD = 9.24, p = .036). This indicated that while chronic CORT, alone or in combination with repeated ketamine or mPFC DBS, did not induce a deficit in locomotion, a combination of all three reduced average velocity.

### 5.2 Changes in mPFC oscillatory activity immediately after treatment

To investigate the acute oscillatory effects of the treatments, we first visualized the power spectral density plots immediately before and after treatment, as well as the percent change between them, in Figure 3A-C. We then compared percent changes in sample entropy and the relative power of the delta, theta, alpha, beta, low gamma, high gamma, and HFO frequency bands between groups immediately after treatment on PSD 24 and 29. Additionally, we tested whether percent changes in sample entropy and the relative power of each frequency band differed significantly from their baseline values. Furthermore, to explore differences in these changes after repeated treatment administration, we compared the percent changes in relative power and sample entropy between the first treatment day, PSD 24, to the sixth treatment day, PSD 29. These comparisons are visualized in Figure 4. Only a subset of animals had valid recordings on each day (PSD 24 and 29: n = 39).

**Figure 3.**
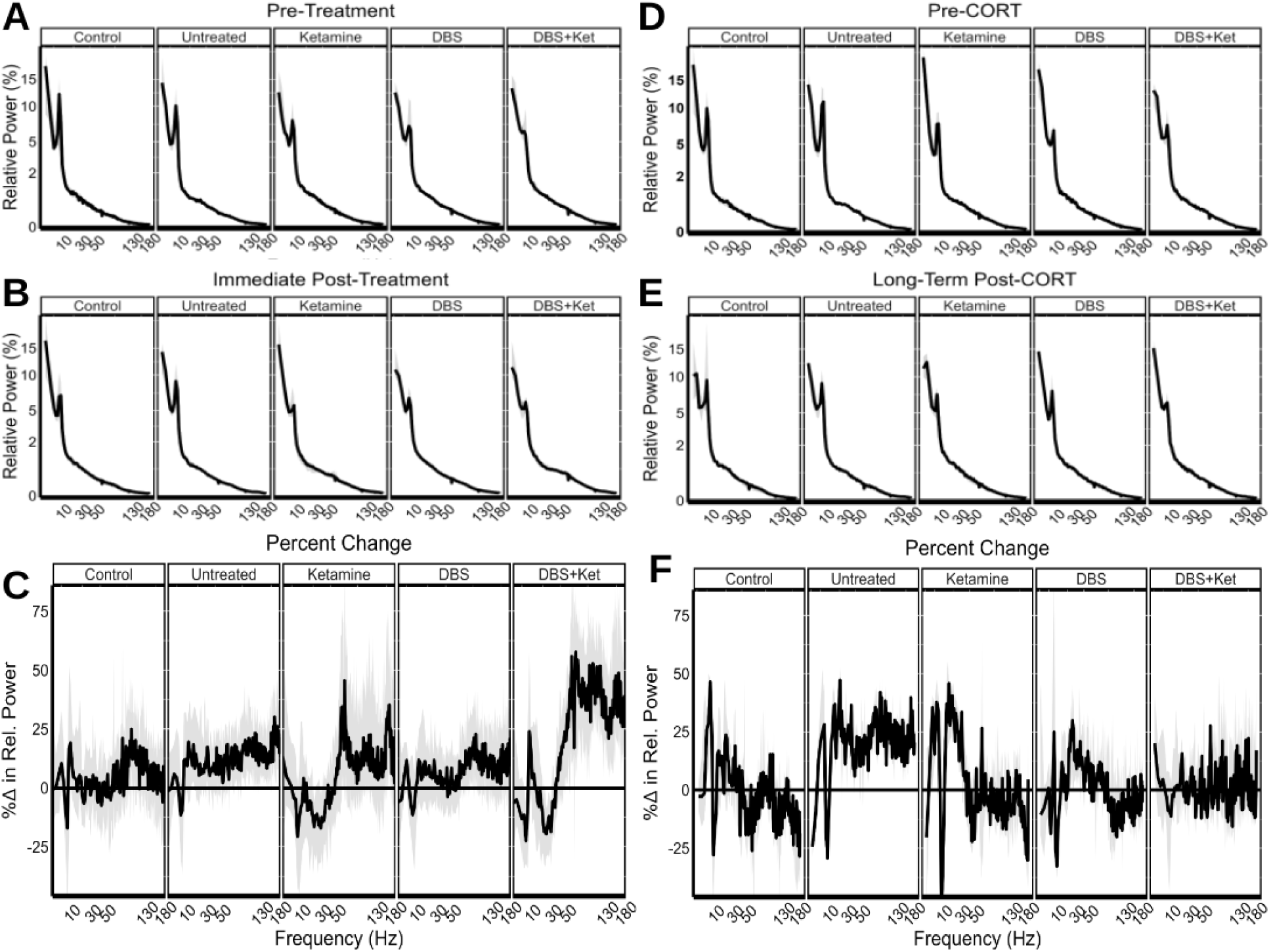
Changes in power spectral density (PSD) plots. PSD plots, with square root transformed axes for improved visualization, of local field potential recordings taken immediately pre-(A) and post-treatment (B), averaged over two treatment days. C, Percent change between A and B, with the x-axis square root transformed. Similar PSD plots from the pre-CORT baseline (D) and long-term post-CORT timepoint (E), with percent change calculated below (F). (CORT = corticosterone)

**Figure 4.**
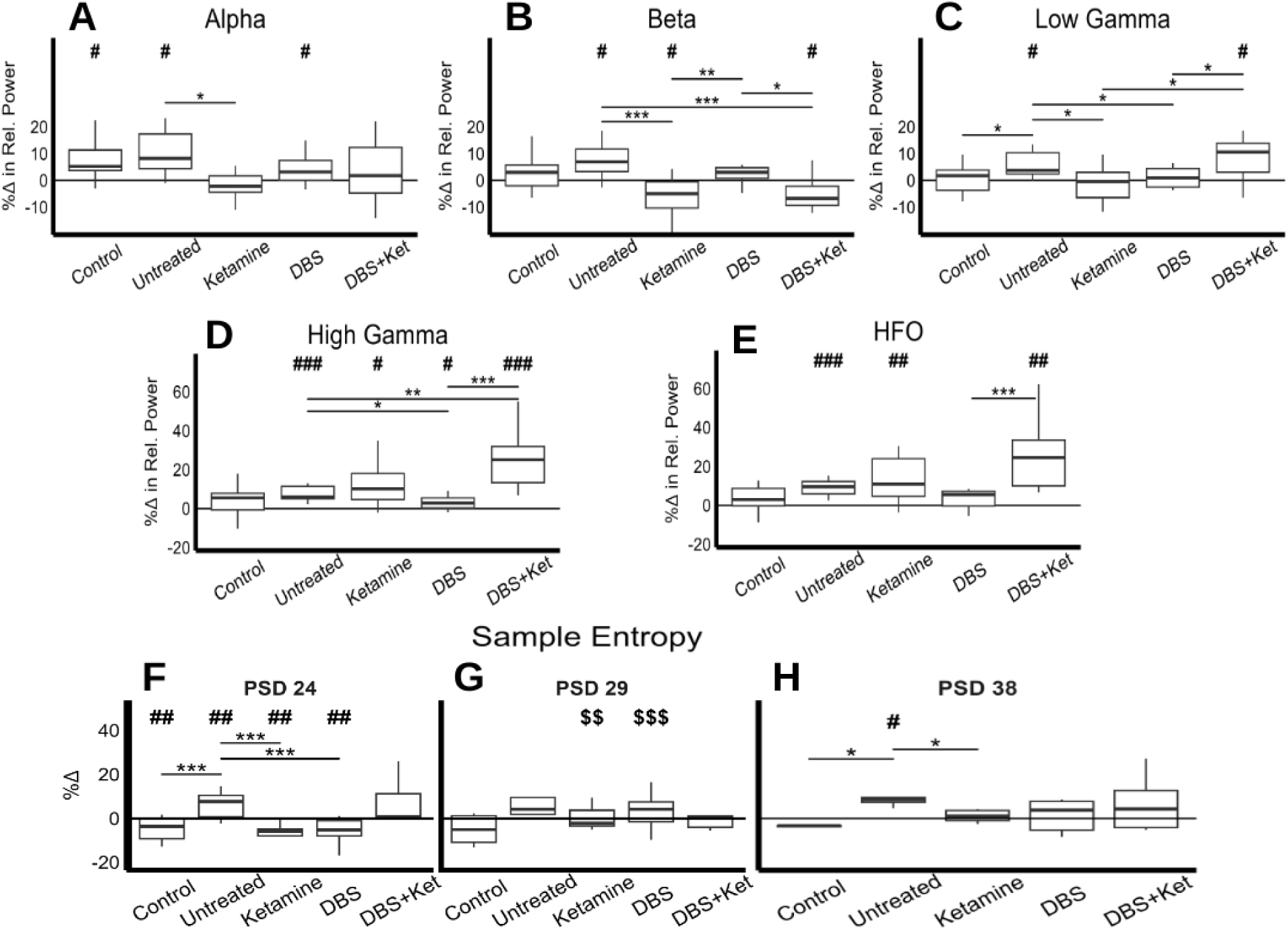
Changes in local field potential (LFP) measures. A-E, Boxplots of LFP oscillatory measures that were significantly altered immediately after treatment, averaged across two treatment days. F-G, Boxplots of the percent changes in sample entropy immediately after treatment on two treatment days. H, Boxplot of the percent changes in sample entropy one week after corticosterone (CORT) and treatment were offset, compared to a pre-CORT baseline. Results of post-hoc tests between groups are shown with stars (*), results of one-sample tests comparing group averages to their pre-CORT or -treatment baseline are shown with number signs (#), and results of one-sample tests comparing group averages between treatment days are shown with dollar signs ($). (*, #, $: p < .05; **, ##, $$: p < .01; ***, ###, $$$: p < .001)

A repeated measures two-way ANOVA with group and day effects was performed on the percent change in relative power for each frequency band. This revealed a significant main effect by group in alpha (F(4,1) = [3.72], p = .018), beta (F(4,1) = [5.95], p < .01), high gamma (F(4,1) = [5.93], p < .01), and HFO (F(4,1) = [6.65], p < .01, all FDR corrected). The group effects in low gamma (F(4, 1) = [2.63], p = .081) showed trends strong enough to warrant further analysis. No significant fixed effects were found by day, and there were no significant group x day interactions for any frequency band, therefore, the days were averaged before post-hoc analysis. The lack of within-group differences between PSD 24 and PSD 29 indicates that treatment-induced changes in relative power in the mPFC were not altered after repeated administration of either ketamine or mPFC DBS.

Similarly, for the percent change in sample entropy, a two-way ANOVA found main group effects (F(4,1) = 10.1, p < .0001). In contrast to the power findings, there was a significant group x day interaction (F(4,1) = [4.48], p < .01), and therefore PSD 24 and 29 were not averaged before post-hoc test. These within-group differences indicate that treatment-induced changes in sample entropy of the mPFC are altered after repeated administration, in contrast to oscillatory activity changes.

#### 5.2.1 Relative alpha power increased in the control, untreated, and DBS groups, but not the ketamine or DBS+ketamine groups

Relative alpha power increased compared to baseline in the control (M = 8.98%, SD = 10.5, p = .045), untreated (M = 10.1%, SD = 8.73, p = .026), and DBS (M = 3.85%, SD = 5.31, p = .048) groups (Fig. 4A). The untreated group differed significantly from the ketamine group (M = −0.89%, SD = 7.79, p = .030), but not the DBS+ketamine group (M = 3.18%, SD = 10.9, all FDR corrected).

#### 5.2.2 Relative beta power increased in the untreated group but decreased in ketamine and DBS+ketamine groups

Compared to baseline, relative beta power increased in the untreated group (M = 7.61%, SD = 6.40, p = .02), but decreased in the ketamine (M = −6.58%, SD = 8.69, p = .035) and DBS+ketamine (M = −4.76%, SD = 6.10, p = .045) groups (Fig. 4B). While relative beta power did not increase in either the control (M = 2.57%, SD = 6.23) or DBS (M = 3.14%, SD = 6.87) groups, they did not significantly differ from the untreated group. In contrast, the ketamine and DBS+ketamine groups differed from the untreated (Ketamine: p < .001; DBS+Ket: p < .001) and DBS groups (Ketamine: p < .01; DBS+Ket: p = .019; all FDR corrected).

#### 5.2.3 Relative low gamma power increased in the untreated and DBS+ketamine groups

Relative low gamma power increased in comparison with baseline in the untreated (M = 5.80%, SD = 4.89, p = .023) and DBS+ketamine (M = 10.2%, SD = 11.0, p = .030) groups (Fig. 4C). The changes in these groups differed from the control (M = .46%, SD = 5.83, untreated: p = .035), ketamine (M = −0.53%, SD = 8.23, untreated: p = .041; DBS+Ket: p = .041), and DBS (M = 1.02%, SD = 3.74, untreated: p = .041; DBS+Ket: p = .041, all FDR corrected) groups, but not from each other.

#### 5.2.4 DBS reduced the increase in relative high gamma power observed in the untreated group, while DBS+ketamine potentiated it

All groups except the control group (M = 3.99%, SD = 8.19) increased in relative high gamma power compared to baseline (untreated: M = 7.52%, SD = 3.83, p < .001; ketamine: M = 12.5%, SD = 17.6, p = .025; DBS: M = 3.17%, SD = 3.39, p = .013; DBS+Ket: M = 24.4%, SD = 14.0, p < .001) (Fig. 4D). Relative high gamma power in the untreated group increased more than in the DBS group (p = .028), and it increased more in the DBS+ketamine group than in the untreated (p < .01) and DBS group (p < .0001, all FDR corrected). While the control group did not increase in relative high gamma power, it did not statistically differ from the untreated group.

#### 5.2.5 HFO power increased in the untreated, ketamine, and DBS+ketamine groups

Relative HFO power increased in the untreated (M = 8.91%, SD = 4.39, p < .001), ketamine (M = 13.2%, SD = 12.1, p < .01), and DBS+ketamine groups (M = 24.3%, SD = 16.6, p < .01) (Fig. 4E). However, the only significant difference amongst the groups was between the DBS+ketamine and DBS groups (M = 4.25%, SD = 6.98, p < .0001, all FDR corrected).

#### 5.2.6 Sample entropy increased in the untreated group, while it decreased in the control, DBS, and ketamine-only groups

On the first day of treatment (PSD 24), sample entropy increased in the untreated group (M = 5.24%, SD = 6.85, p < .01), and decreased in the control (M = −5.02%, SD = 6.54, p < .01), ketamine (M = −8.00%, SD = 9.82, p < .01), and DBS (M = −6.48%, SD = 7.02, p < .01) groups (Fig. 4F). While sample entropy in the DBS+ketamine group (M = 5.16%, SD = 15.9) did not change, it did not statistically differ from the untreated group. In contrast, the control (p < .0001), ketamine (p < .0001), and DBS (p < .0001, all FDR corrected) groups differed from the untreated group.

#### 5.2.7 After repeated administration, DBS and ketamine no longer reduced sample entropy from baseline; however, ketamine and DBS+ketamine inhibited the increase found in the untreated group

By the sixth day of treatment (PSD 29), sample entropy did not change in the DBS (M = 3.38%, SD = 9.11) or ketamine (M = −0.02%, SD = 5.95) groups relative to baseline, and the DBS group did not differ from the untreated group (M = 10.4%, SD = 14.0) (Fig. 4G). Both groups significantly differed from their changes on the first day of treatment (DBS: p < .01; ketamine: p < .001). However, like on PSD 24, ketamine had significantly lower sample entropy than the untreated group (p < .001), where sample entropy increased significantly (p < .01). Interestingly, in contrast to the first treatment day, sample entropy on PSD 29 was significantly lower in the DBS+ketamine group (M = −2.48%, SD = 13.2) than in the untreated group (p < .01), though compared to the first day the response of the DBS+ketamine group did not statistically change. Finally, as on PSD 24, the untreated group differed from the control group (M = −4.82%, SD = 6.84, p < .0001, all FDR corrected), in which sample entropy decreased (p = .026).

### 5.3 Changes in mPFC oscillatory activity one day (short-term) and one week (long-term) after chronic CORT and treatment

To explore the short- and long-term effects of CORT and treatment on LFP spectral power and sample entropy in the mPFC, we compared the relative spectral powers and sample entropies of the groups at two timepoints after CORT and treatment offset, PSD 31 and PSD 38, to the timepoint before the first CORT injection, PSD 10. Only a subset of animals had valid before and after LFP recordings (PSD 31: n = 33; PSD 38: n = 31).

A one-way ANOVA comparing groups was performed on these percent changes. For spectral power, no frequency bands differed significantly on either day after FDR correction. The power spectral density plots at the pre-CORT and long-term post-CORT timepoint, PSD 38, as well as the percent change between them, are visualized in Figure 3D-F. For sample entropy, at the long-term timepoint, PSD 38, the ANOVA showed trends strong enough to warrant further investigation (F[4,26] = 2.63, p = .057), and post-hoc tests yielded robust differences. The untreated group (M = 8.64%, SD = 3.13) had increased sample entropy (p = .018) relative to pre-CORT baseline, and differed from the control (M = −5.75%, SD = 6.90, p = .016) and ketamine (M = −0.61%, SD = 6.68, p = .010, all FDR corrected) groups, but not the DBS (M = 1.43%, SD = 7.71) and DBS+ketamine (M = 6.43%, SD = 12.0) groups (Fig. 4H).

### 5.4 Correlates of altered behavior on the Groom Test and the Forced Swim Test

To investigate the hypothesis that long-term electrophysiological changes in the mPFC relate to depression-like symptoms and their rescue, we correlated evoked groom time in the GT, immobility time in the FST, and relative weight on PSD 38 with the altered electrophysiological variable on PSD 38, sample entropy. Correlation coefficients and p-values were calculated for the control and ketamine groups, each of which were pooled with the untreated group. We found that the percent change in sample entropy by PSD 38 correlated with evoked groom time in the untreated x ketamine group pool (p = .030, r = .-63), as shown in Figure 5A. Similarly, to examine the hypothesis that cognitive improvements related to the rescue of depression-like symptoms, we correlated the pooled untreated x DBS group NORT scores with evoked groom and immobility time. Indeed, the percent change in discrimination index correlated with evoked groom time (p = .019, r = .58) (Fig. 5B).

**Figure 5.**
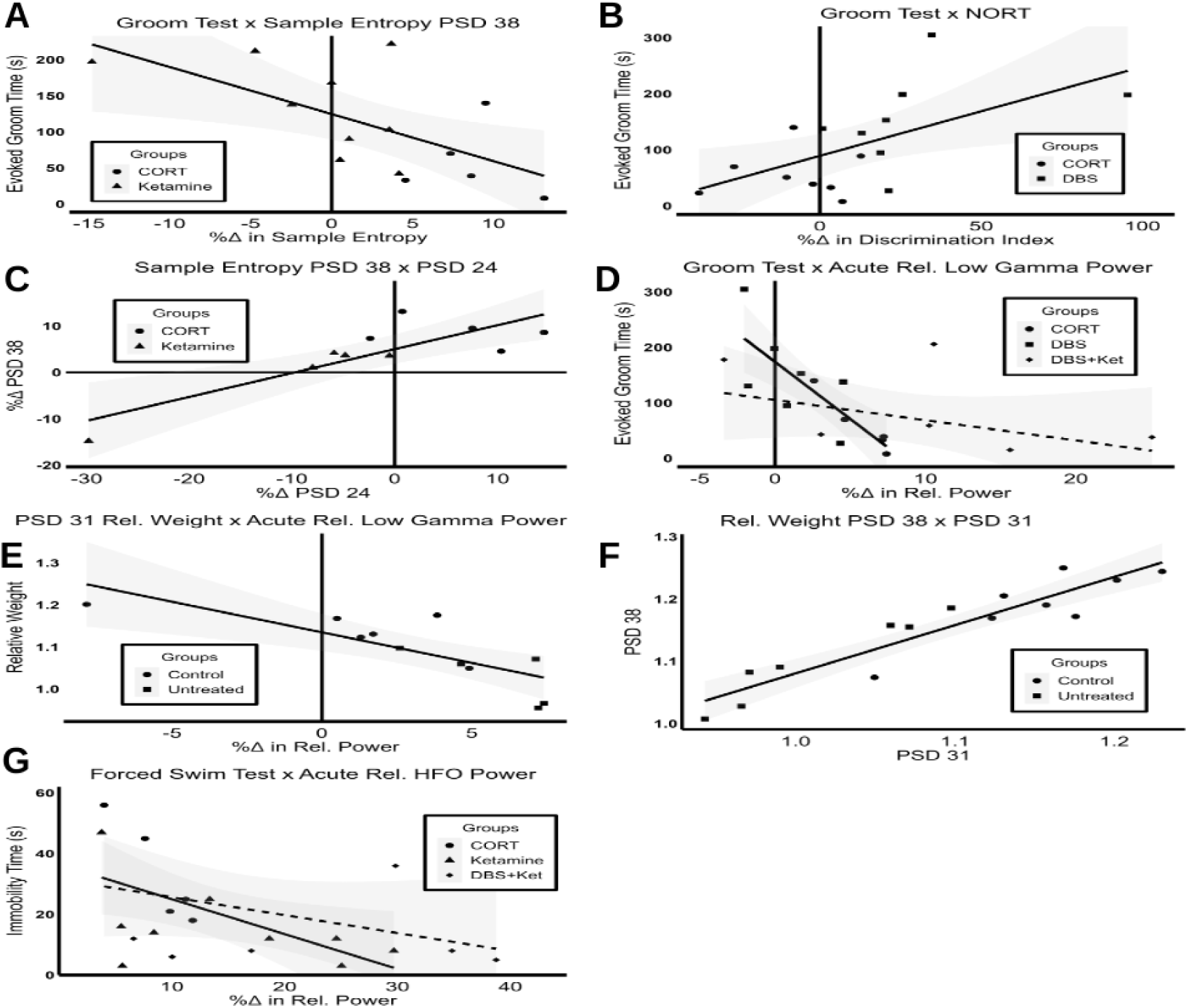
Correlations among electrophysiology measures and depression-like symptoms. A, In the untreated x ketamine group pool, the percent change in sample entropy from pre-CORT baseline and the long-term post-CORT timepoint, PSD 38, correlated with evoked groom time in the GT, a measure of apathy-like behavior. B, In the untreated x DBS group pool, evoked groom time in the GT was correlated with the percent change in DI in the NORT, a measure of object memory. C, In the untreated x ketamine group pool, the percent change in long-term sample entropy on PSD 38 correlated with the percent change immediately after treatment on PSD 24. D, In the untreated x DBS group and untreated x DBS+ketamine group pools, the percent change in relative low gamma power immediately after treatment correlated with evoked groom time in the GT. E, In the untreated x control group pool, the percent change in relative low gamma power immediately after treatment correlated with relative weight on PSD 30, which correlated with relative weight on PSD 38 (F). G, In the untreated x ketamine group pool, the percent change in relative HFO power immediately after treatment correlated with immobility time in the FST, a measure of despair-like behavior. (CORT = corticosterone, PSD = post surgery day, DBS = deep brain stimulation, GT = Groom Test, DI = Discrimination Index, NORT = Novel Object Recognition Test, HFO = high frequency oscillation)

As we had found long-term behavioral and electrophysiological changes that were correlated with the rescue of the GT and FST, we tested whether they related to electrophysiological changes immediately after treatment. Specifically, in the untreated x ketamine group pool, acutely altered LFP measures were correlated with sample entropy on PSD 38, and in the untreated x DBS group pool they were correlated with NORT performance. For the untreated x ketamine group pool, we found that the decrease in sample entropy immediately after treatment on PSD 24 correlated with the decreased sample entropy on PSD 38 (p < .01, r = .84) (Fig. 5C). Interestingly, relative low gamma power at this time did not correlate with the NORT outcome in the untreated x DBS group pool.

Then, to explore the hypothesis that acute changes in electrophysiological measures immediately after treatment related directly to long-term behavioral and weight measures, we correlated the acutely altered measures of each group (pooled with the untreated group) with evoked groom time, immobility time, and weight on PSD 30 and 38. In the untreated x DBS group pool, relative low gamma power immediately after treatment negatively correlated with evoked groom time (p < .01, r = -.82), a finding that was replicated in the untreated x DBS+ketamine group pool (p = .014, r = -.97) (Fig. 5D). Meanwhile, in the untreated x control group pool, we found that the depression-like deficit in weight gain on PSD 30 and 38 correlated with relative low gamma power immediately after sham treatment (PSD 30: p < .01, r = -.86; PSD 38: p < .01, r = -.90) (Fig. 5E), and that the weights on these days correlated with one another (p < .0001, r = .93) (Fig. 5F). Finally, there was a strong trend in the untreated x ketamine group pool where the increase in relative HFO power immediately after treatment correlated with decreased immobility time in the FST (p = .054, r = .59), a correlation which was significant in the untreated x DBS+ketamine group pool (p = .048, r = −0.95, all FDR corrected) (Fig. 5G).

Finally, to ensure that differences in weight and locomotion did not confound our measures of depression-like behavior, we correlated each pool’s relative weights on PSD 38, and their average velocity in the OFT, with evoked groom and immobility time. We found no significant correlations, evidence that the deficits observed in the GT and FST were not due to non-specific motor or metabolic deficits.

## 6. Discussion

### 6.1 Changes in mPFC LFP measures did not correlate with depression-like behavior, but did correlate with weight gain deficits

Chronic CORT induced the primary depression symptoms, apathy and despair, in the Groom and Forced Swim Tests, respectively, as well as a secondary symptom, a deficit in weight gain (Kennedy, 2008) (Fig. 2). It is thought that chronic CORT induces a depression-like phenotype through disruptions in the acute stress-response activation of structures responsible for emotional and cognitive functions, like the mPFC, such that their long-term function is disrupted days to weeks post-offset (Sterner & Kalynchuk, 2010). The mPFC is particularly impacted in this process (Pizzagalli & Roberts, 2022) due to a relatively high density of CORT-sensitive receptors (McEwen et al., 1986). Long-term electrophysiological changes here are highly implicated in both MDD (Fingelkurts & Fingelkurts, 2015; Knott et al., 2001; Mumtaz et al., 2017) and its preclinical models (Bühning et al., 2022; Claverie et al., 2016; Zheng et al., 2012), and therefore have gained attention as etiological and therapeutic biomarkers (Fitzgerald 2018). However, the acute changes in mPFC activity that represent the aberrant, depression-inducing stress-response have not been characterized. Therefore, we compared the LFP recordings of the control and untreated groups immediately after the mild stress of sham treatment during the final week of depression induction (Fig 1A). Indeed, there were significant increases in sample entropy and relative low gamma in the untreated group relative to the control group (Fig. 4) that may have represented aberrant stress-induced hyperactivity. Both have been associated with levels of neural activity and function (Fitzgerald & Watson, 2018; Wang et al., 2018), as well as stress-responses (Bercum et al., 2021; Tran et al., 2007), however, their functional significance is unclear.

Furthermore, this study was also designed to discover long-term electrophysiological changes after chronic CORT that related to long-term behavioral deficits. Interestingly, in the LFP recording that was taken over a week after the offset of CORT, sample entropy had decreased in the control group yet increased in the untreated group compared to their pre-CORT baselines (Fig. 4). While this recording on PSD 38 did not occur on the same day as the GT (PSD 35) or the FST (PSD 39), the depression-like state induced by chronic CORT, and its electrophysiological correlates, are relatively stable once induced (Sterner & Kalynchuk, 2010). Therefore we believe any changes observed in this recording represent long-term changes in mPFC activity that are sustained across the timeframe of the behavioral assay. While the functional significance of sample entropy is currently unknown, changes in this measure have been observed in depression before, though clinical studies disagree regarding whether frontal sample entropy increases or decreases (Acharya et al., 2015; Čukić et al., 2020; Faust et al., 2014; Lin et al., 2020). Similarly, a previous preclinical study found decreases in the mPFC after chronic unpredictable stress (Zheng et al., 2012), while the present study found the opposite. The differences in findings likely relate to differences in methodology. For example, while the depression model employed by Zheng and colleagues relied on the same mechanisms as those in the present study, and they used similar parameters for calculating sample entropy, there were important differences in LFP sample rate and referencing. Despite inconsistent methodologies, the repeated finding that sample entropy differs between control, disease, and treatment groups, in both humans and animals, supports its potential as a depressive biomarker.

To further characterize the relationships between mPFC activity measures and depression-like symptoms, and thereby further evaluate their potential as biomarkers, we correlated them with immobility time in the FST, evoked groom time in the GT, and relative weight at short- and long-term timepoints post-CORT. While no electrophysiological changes correlated with the depression-like behavioral measures, relative low gamma power immediately after treatment negatively correlated with relative weight on both PSD 31 and PSD 38 (Fig. 5). The pathophysiology of weight loss after chronic corticosterone is not understood, however, we hypothesize that increases in low gamma power represented a functional change that altered the mPFC’s role in homeostatic food-related decision-making (Eiselt et al., 2021), as well as an overactivation in neural activity as part of a larger aberrant stress-response that included metabolic changes (Karatsoreos et al., 2010; Whittington et al., 2011). While the mechanism requires further study, this data suggests that the acute changes in relative low gamma power response during stress-based disease induction could be a predictive biomarker for future weight symptoms. In addition, while low gamma and sample entropy did not correlate with depression-like behavior, the significant difference in these measures between the control and depression model animals supports their potential role as biomarkers of depression.

### 6.2 Separately, DBS and ketamine rescued depression-like behaviors, which correlated with contemporary long-term changes in mPFC LFP and cognition measures

After confirming that CORT induced depression-like symptoms, we validated the therapeutic effectiveness of two treatment modalities, repeated mPFC DBS and ketamine. Separately, each rescued behavioral performance in tests of the primary symptoms of depression, apathy (GT) and despair (FST) (Fig. 2). Interestingly, neither modality rescued a secondary depression symptom induced by CORT, a deficit in weight gain (Fig. 2). We tested the hypothesis that long-term changes in mPFC LFP and improvements in cognitive function correlated with the observed antidepressant effects on immobility and evoked groom time. Only variables that differed from the untreated group could explain differences in these behaviors, therefore sample entropy on PSD 38 was considered for the untreated x ketamine group pool, and the NORT was investigated in the untreated x DBS group pool. We found that the long-term normalization of sample entropy in the ketamine pool significantly correlated with the antidepressant effect on apathy-like symptoms in the GT (Fig. 5C). While the functional significance of sample entropy in neurophysiology is unclear, it is thought to represent information processing within a region, and therefore its degree of functional activity and connectivity (Wang et al., 2018). Interestingly, dysfunctional emotion regulation by the mPFC is a well-characterized driver of apathy in depression (Levy & Dubois, 2006), and human fMRI studies have shown that ketamine induces sustained normalization of activity in emotion processing structures following valenced stimuli (Reed et al., 2018). Therefore, we hypothesize that ketamine’s long-term modulation of sample entropy related to a normalization of emotion regulation within the mPFC, which rescued apathy-like behavior. However, given that the strength of the correlation was moderate (Fig. 5C), there were likely other regions or functional changes which related to the rescue of this behavior. In addition, we cannot determine causality from these correlations, however, they suggest that sample entropy could function as a biomarker of ketamine’s efficacy in treating apathy-like symptoms. Finally, the long-term normalization of sample entropy by ketamine did not correlate with the rescue of the despair-like symptoms in the FST, therefore long-term changes outside of those studied here must have driven the rescue of immobility time.

Meanwhile, in the DBS group, we found that the improvement in recognition memory in the NORT correlated with the rescue of apathy-like symptoms in the GT (Fig. 5A). In addition to the aforementioned emotional deficits, cognitive deficits stemming from mPFC dysfunction have also been implicated in the etiology of apathy (Levy & Dubois, 2006). Particularly implicated are two components of recognition memory, working memory and episodic memory, which involve the mPFC (Blumenfeld & Ranganath, 2007; Ragozzino et al., 2002), and are measured by the NORT (Antunes & Biala, 2012). Furthermore, working memory has been previously correlated with apathy in human schizophrenia patients (Raffard et al., 2016). We therefore hypothesize that part of the therapeutic mechanism of mPFC DBS in the alleviation of apathy-like symptoms was the improvement of these two aspects of cognitive function. It is important to note that, in addition to causality, directionality can not be established from these correlations, such that decreases in apathy perhaps underpinned increased interest in novel objects. The improvement in NORT, and therefore the putative improvements in working memory and episodic memory, after DBS did not relate the improvement in the FST. Thus, like in the ketamine group, functional changes outside of those we examined in the mPFC may have underpinned the improvement in despair-like symptoms.

### 6.3 Changes in mPFC LFP immediately after treatment correlated with the long-term antidepressant behavioral effects of DBS and ketamine, but not necessarily their long-term electrophysiological and cognitive effects

After determining which long-term improvements in mPFC and cognitive function correlated with contemporary improvements in depression-like behaviors, we then explored which immediate effects correlated with these long-term changes. These relationships could potentially bridge the gap between the more commonly studied acute effects of these treatment modalities, and their relatively understudied sustained effects. Again, only electrophysiological measures that were significantly altered were tested. First, we found that the benefit of ketamine in FST immobility time nearly correlated with an immediate increase in relative HFO power (Fig. 5G). Increases in HFO power after NMDA receptor antagonist administration are well documented, and likely relate to the fast-spiking GABAergic interneurons they depend on (Jackson et al., 2011). The GABAergic interneurons of the mPFC are known to be dysfunctional in depression (Duman et al., 2019), therefore this acute HFO modulation may represent a stimulus for long-term neurophysiological changes that have been implicated in the therapeutic action of ketamine, particularly synaptic plasticity and potentiation (Cornwell et al., 2012; Duman et al., 2019). As subanesthetic ketamine preferentially antagonizes the activity of NMDA receptors in GABAergic interneurons, it induces net excitation and positively modulates synaptic plasticity and potentiation through mechanism such as brain derived neurotrophic factor (BDNF) (Duman et al., 2019) and synaptic remodeling (Sala et al., 2022). Increased relative HFO activity after ketamine treatment may therefore function as a therapeutic biomarker. However, as this correlation was only a trend in the ketamine group, and relative HFO power also increased in the untreated group, its potential utility as a biomarker for treatment’s mechanism and efficacy will require further study. We hypothesize that the HFO activity observed in the untreated group merely reflected spike contamination due to aberrant increases in firing rates due to stress (Wilber et al., 2011). Meanwhile, there is evidence that HFO activity in the ketamine treated groups was theta-associated HFO, which has been observed after ketamine (Wróbel et al., 2020), and has been hypothesized to play a functional role during REM sleep (Scheffer-Teixeira et al., 2013) and fear extinction (Watanabe et al., 2021). However, since no long-term measures in the ketamine group correlated with immobility time, the functional changes through which these acute changes in relative HFO power improved FST performance is also unclear. In contrast, in terms of the rescue of apathy-like symptoms in the GT, reduced sample entropy on the first day of ketamine treatment correlated with reduced sample entropy in the long-term. Like the modulation of long-term sample entropy, we hypothesize that this acute normalization relates to the normalization of mPFC emotion processing activity after ketamine treatment. Further, based on our correlations, we posit that the immediate normalization of those processes drove their long-term normalization, though future experiments are needed to demonstrate a causal effect of sample entropy. The present study indicates that both acute and long-term measurements of sample entropy in the mPFC can be used as biomarkers of treatment efficacy regarding apathy-like symptoms after ketamine treatment.

Meanwhile, in the mPFC DBS group, we found that suppression of relative low gamma power immediately after treatment correlated with improvement in the GT (Fig. 5B). Low gamma power is generated via interplay between pyramidal glutamatergic neurons and GABAergic interneurons (Whittington et al., 2011), precisely the cell types that experience excitotoxicity-induced atrophy and dysfunction due to stress (Duman et al., 2019). We hypothesize that DBS protected the structure and function of these mPFC cells by inhibiting their hyperactivity during depression induction (Etiévant et al., 2015; Sprengers et al., 2020), and thereby inhibiting the development of apathy-like symptoms (Zamboni et al., 2008). This finding parallels results in humans, in which the suppression of low gamma in the mPFC immediately after DBS correlated with decreased depressive symptoms (Sun et al., 2015). However, interestingly, the acute changes in relative low gamma power did not correlate with the improvements in the NORT, which had correlated with the GT. Therefore, it remains unclear how acute suppression of low gamma power translated to the long-term improvements in the GT. However, measurements of low gamma power immediately after mPFC DBS may function as biomarkers for treatment efficacy, in terms of apathy-like symptoms. Finally, no altered electrophysiological or behavioral changes correlated with the rescue of the FST after mPFC DBS, therefore other regions or electrophysiological changes must have underpinned its rescue of despair-like symptoms.

### 6.4 Combined, DBS and ketamine did not fully rescue the GT, supporting the therapeutic importance of certain electrophysiological and cognitive correlates of antidepressant behavioral effects

Finally, after finding correlates of rescued behavior in both treatment modalities separately, we hypothesized that a novel combinatorial treatment would be synergistically effective. We found that this approach fully rescued despair-like symptoms in the FST, however, it failed to fully rescue apathy-like symptoms in the GT (Fig. 2A, B). As when each treatment was administered separately, the combinatorial treatment also did not rescue the depression-like deficit in weight gain induced by chronic CORT (Fig. 2G, H). To understand why our synergistic hypothesis was refuted, we analyzed immediate and long-term mPFC LFP changes to find correlates of therapeutic successes and failures. First, we found that a lack of suppression in low gamma power immediately after treatment correlated with the DBS+ketamine group’s apathy-like symptoms in the GT (Fig. 5B). This was in contrast to the DBS group, where a suppression of low gamma power correlated with the rescue of evoked groom time (Fig. 5B). In addition, the NORT was not improved by the combinatorial treatment, unlike in the DBS group (Fig. 2C). Similarly, in contrast to the ketamine group, sample entropy was not normalized at acute nor long-term timepoints (Fig. 4F-H) in the combination group. Furthermore, the NORT and sample entropy were no longer correlated with the GT. Taken together, these findings support our conclusions that low gamma power and cognitive function in mPFC DBS, and sample entropy in ketamine, relate to their rescue of apathy-like behavior. Meanwhile, in the FST, increases in relative HFO power correlated with the rescue of despair-like symptoms in the DBS+ketamine group (Fig. 5G). This paralleled the trend observed in the ketamine group, therefore the results of the combination group support our conclusion that an increase in relative HFO power may be important to the therapeutic mechanism of ketamine.

Finally, we observed decreased average velocity in the OFT trials of the combination group, compared to the control group (Fig. 2F). While decreased locomotor activity can occur in depression (American Psychiatric Association, 2013), we found no deficits in the untreated group, therefore this depression-like symptom was unique to the combination group. Importantly, this did not correlate with depression-like behavior, excluding it from confounding the observed apathy-like behavior, and establishing decreased locomotor activity as a unique symptom that emerged from a combination of CORT, mPFC DBS, and ketamine. As it also did not correlate with altered electrophysiology at any timepoint, the underlying neurophysiological change behind this symptom is unclear. However, given that DBS was applied to the mPFC, it likely involved its connections with structures in the corticolimbic circuit governing motor output, such as the striatum, ventral tegmental area and spinal cord (De Haan et al., 2018). The appearance of this symptom after treatment provides further evidence that the combination of mPFC DBS and ketamine, at the dosages examined, was counterproductive in treating depression-like symptoms.

### 6.5 Limitations

There are several important limitations to the present study. Depression differs between sexes in humans (Breslau et al., 1995), yet sex-based differences were not explored here. As the depression-inducing effects of chronic corticosterone are age- and sex-dependent, such that only adult male and adolescent female rats are vulnerable (Bertholomey et al., 2022), adult female rats would not have been etiologically comparable to our cohort of adult male rats. Also, the animals were single housed, which may have altered behavior (Liu et al., 2020). Additionally, our experiment included weekly sucrose preference tests (SPT), however due to faults in the apparatus this test was excluded. For consistency, all animals performed the SPT habituations on PSD 7 and 8, as well as tests on PSD 9, 16, 23, 30, and 37, which had been preceded by 24 hour fasts. Finally, as our LFP recordings were segmented to avoid bouts of noise, there may have been a sampling bias.

### 6.6 Conclusion

This study contributes to the growing evidence that mPFC electrophysiology measures are etiologically and therapeutically relevant functional changes that underlie depression and the antidepressant action of mPFC DBS and ketamine. Therefore, it also supports their utility as predictive biomarkers for depression and treatment efficacy. Specifically, our findings support previous clinical and preclinical studies that associated low gamma power and sample entropy with depression or remission, and could therefore act as translatable, objective biomarkers across treatments and species. Furthermore, our study is among the first to provide evidence that improved aspects of cognition after mPFC DBS, and increased HFO power after ketamine, relate to the therapeutic efficacy of these modalities. Further exploration of the mechanisms that interconnect electrophysiological and behavior changes will enable the development of biomarkers for targeted, personalized, and monitored treatment strategies that can address the weaknesses of current treatments. Finally, the complex interaction between the two treatments studied here enhanced our understanding of their separate mechanisms and demonstrated that the combination of these treatments may be detrimental to certain aspects of therapeutic efficacy. This needs to be explored further, but the nature of their interaction may be critical information for patients and clinicians.

